## Supplementary material for "Dissection of amino acid acquisition pathways in *Borrelia burgdorferi* uncovers unique physiological responses": Spplemental Images and Tables

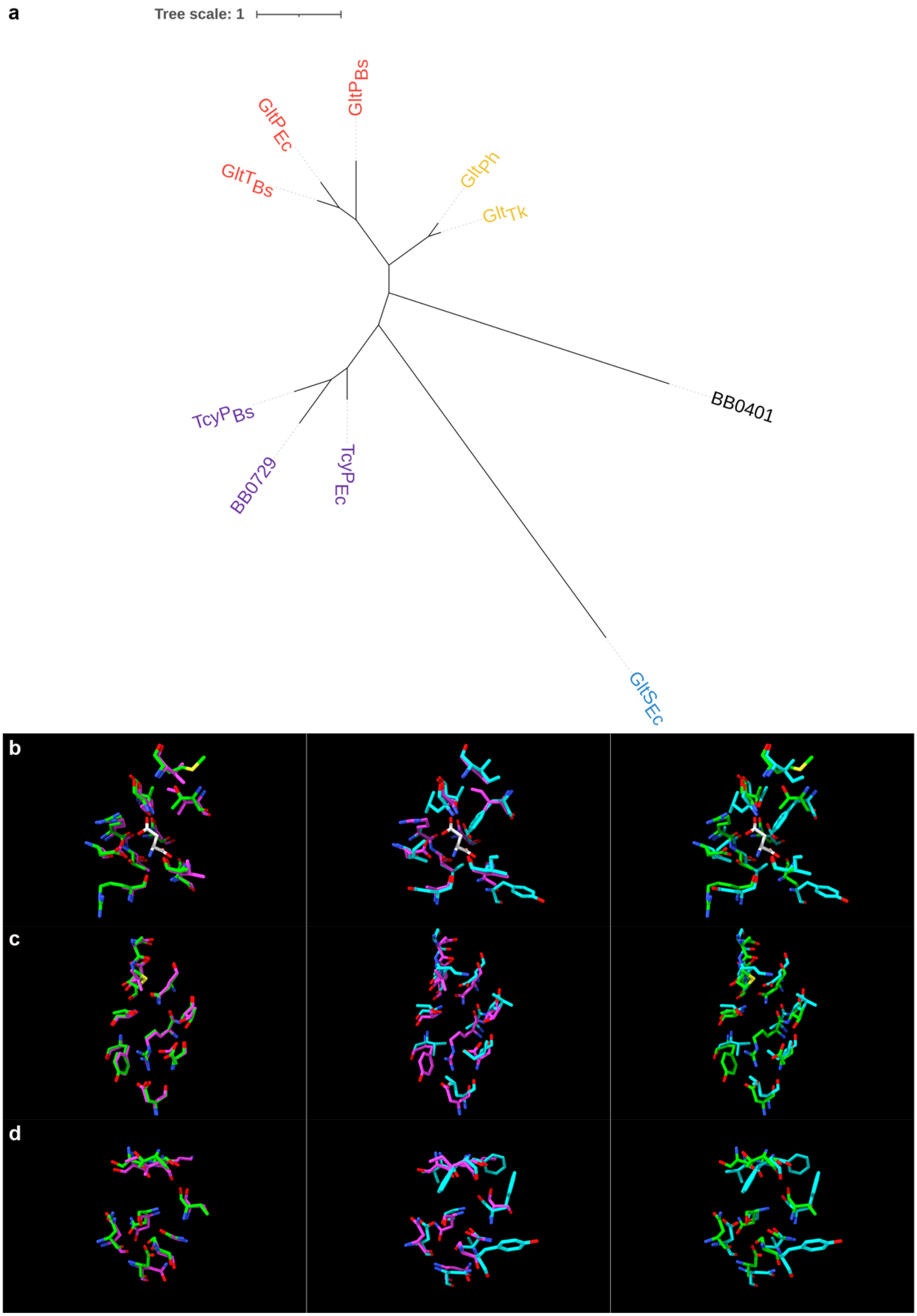


**Fig S1: BB0401 models as a GltP.** a) Unrooted phylogenetic tree of characterized Glt and TcyP transporters. Dedicated glutamate transporters are blue, aspartate transporters are orange, glutamate and aspartate transporters in red, cystine transporters in purple. b-d) Residue alignments for b) aspartate binding site with Asp from 2nwl, c) predicted glutamate binding site, and c) sodium binding sites. $Glt_{Ph}$ (2nwl) is shown in green, $GltP_{Ec}$ model in magenta, and BB0401 model in cyan.

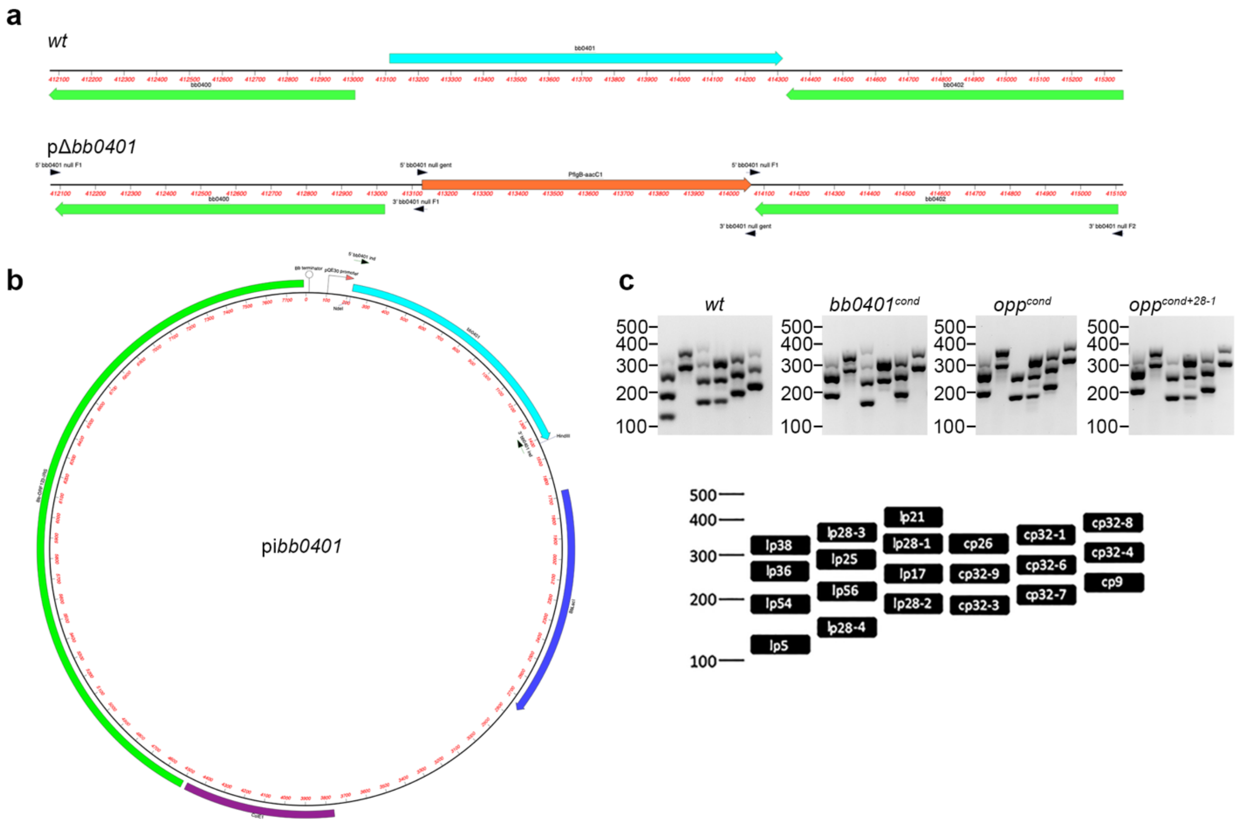


**Fig S2: *bb0401* is essential for growth. a** Schematic of *bb0401* locus on *B. burgdorferi* chromosome (*wt*) where gene of interest is in cyan and construction of pΔ*bb0401* where antibiotic cassette is in orange and arrows represent primer locations **b** Schematic of pi*bb0401* where gene of interest is in cyan, *lacI* in blue, *E. coli* origin of replication in purple, and cp9 shuttle vector region in green, arrows represent primer location, and restriction enzyme sites are shown. **c** Plasmid content multiplex PCRs for all strains, ladder sizes are shown in bp and a schematic of multiplex targets is displayed below.

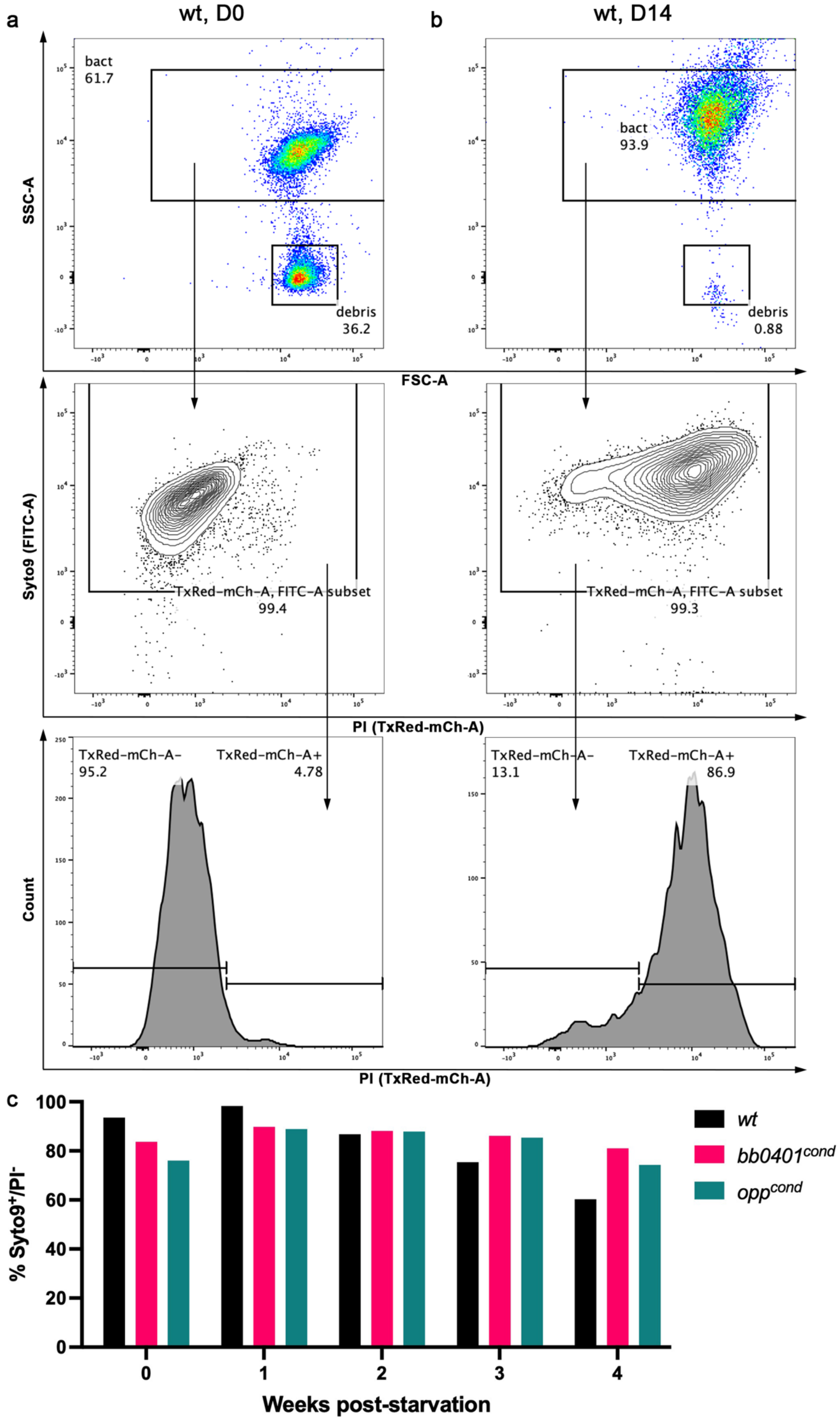


**Figure S3: Amino acid transport is not essential for room temperature growth.** a-b) Example gating strategy to identify live/dead cell populations using flow cytometry for *wt*. Top graph shows FSC-A/SSC-A for exclusion of debris present in the media, middle graph shows TxRed-mCh-A/FITC-A to identify the FITC-A+ subset. Bottom graph shows a histogram of TxRed-mCh-A to gate live and dead populations. c) Percent of live cells (Syto9+/PI-) during room temperature incubation sampled weekly over a four-week period. Two-way ANOVA found no statistical significance in pairwise comparisons.

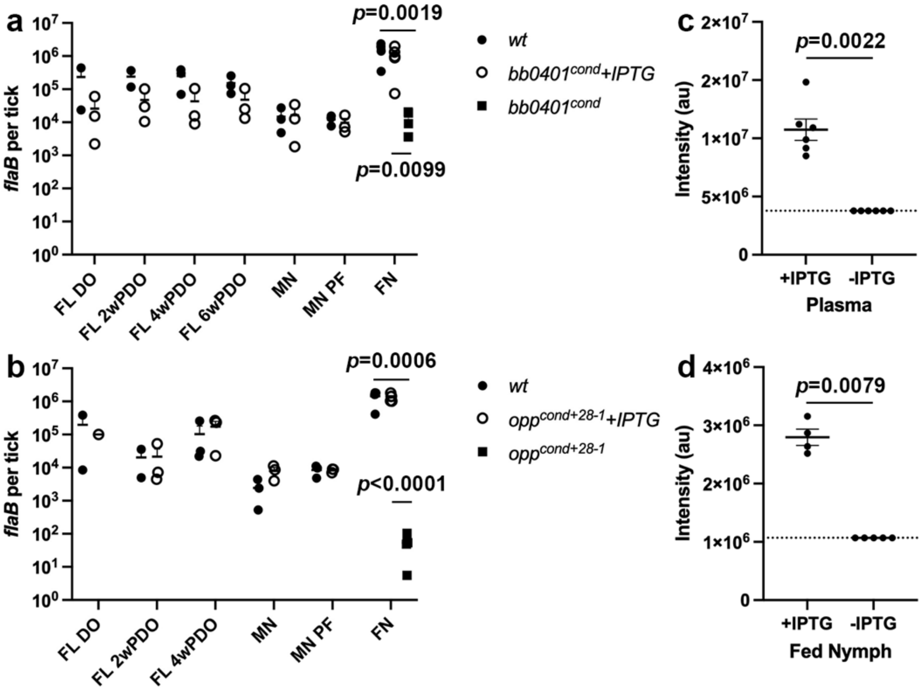


**Figure S4: DNA burdens are lower in fed nymph for both mutants.** Burdens as determined by qPCR of fed larvae at drop-off (FL DO), fed larvae at timepoints post-drop-off (FL #wPDO), post-molt nymphs (MN), molted nymphs prior to feeding (MN PF), and fed nymphs (FN) for a) *bb0401*$^{comp}$ and b) *opp*$^{cond+28\text{-}1}$. IPTG detection in c) pooled mouse plasma and d) pooled fed nymphs. Dotted lined represent the LOQ. *p*-values were determined for pairwise comparisons using a two-tailed unpaired *t* test.

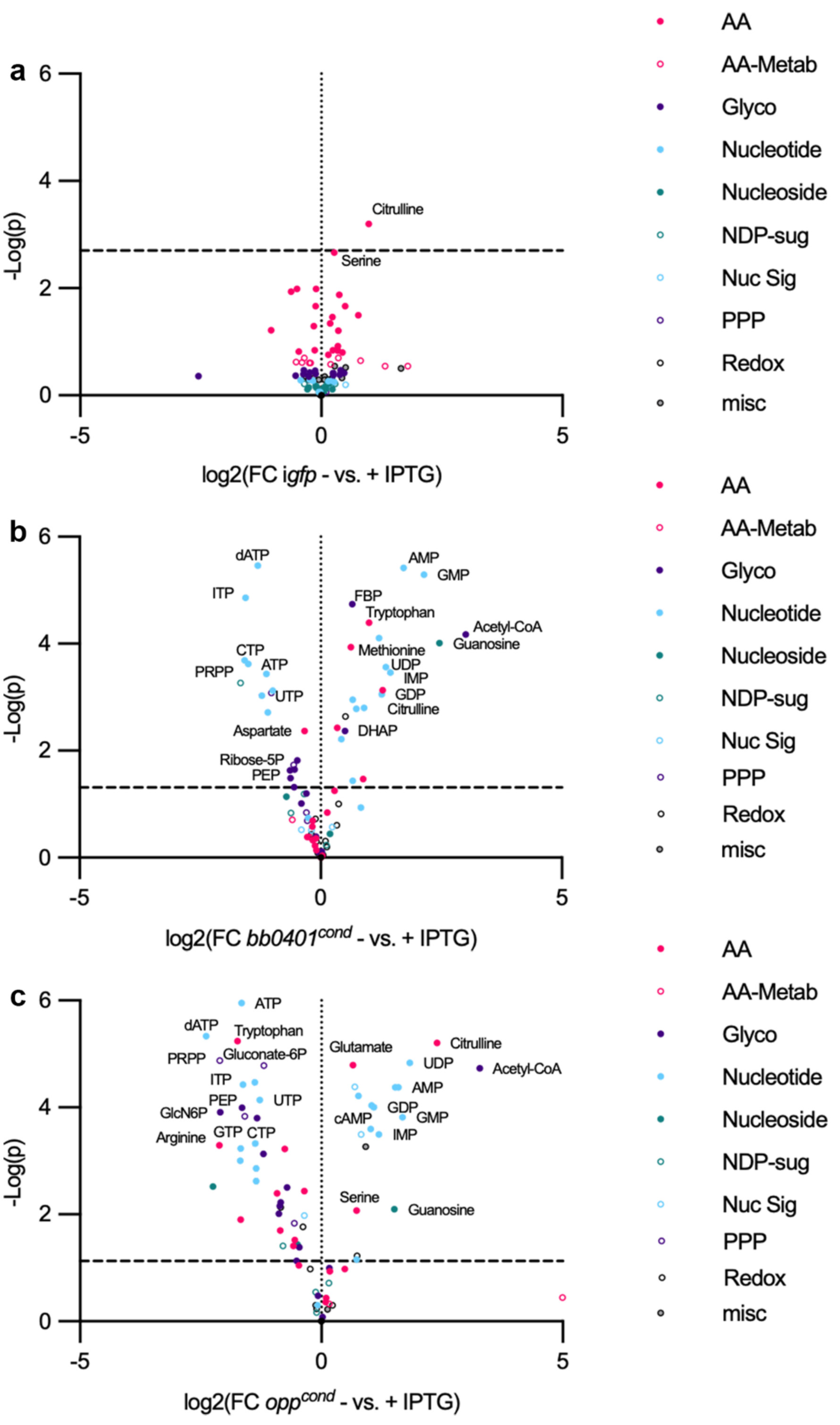


**Figure S5: *opp*$^{cond}$ starvation results is a larger metabolite shift than *bb0401*$^{cond}$.** Volcano plots showing metabolite changes in a) i*gfp* control, b) *bb0401*$^{cond}$, and c) *opp*$^{cond}$ when growth without and with 1 mM IPTG. Metabolites are color-coded by primary pathway. Dotted line represents 10% FDR. Tabulated results can be found in Table S3. AA=amino acids, AA-Metab=amino acid metabolites, Glyco=glycolysis and other carbohydrate, NDP-sug=nucleotide diphosphate sugar conjugates, Nuc Sig=signaling nucelotides, PPP=pentose phosphate pathway, Redox=redox cofactors.

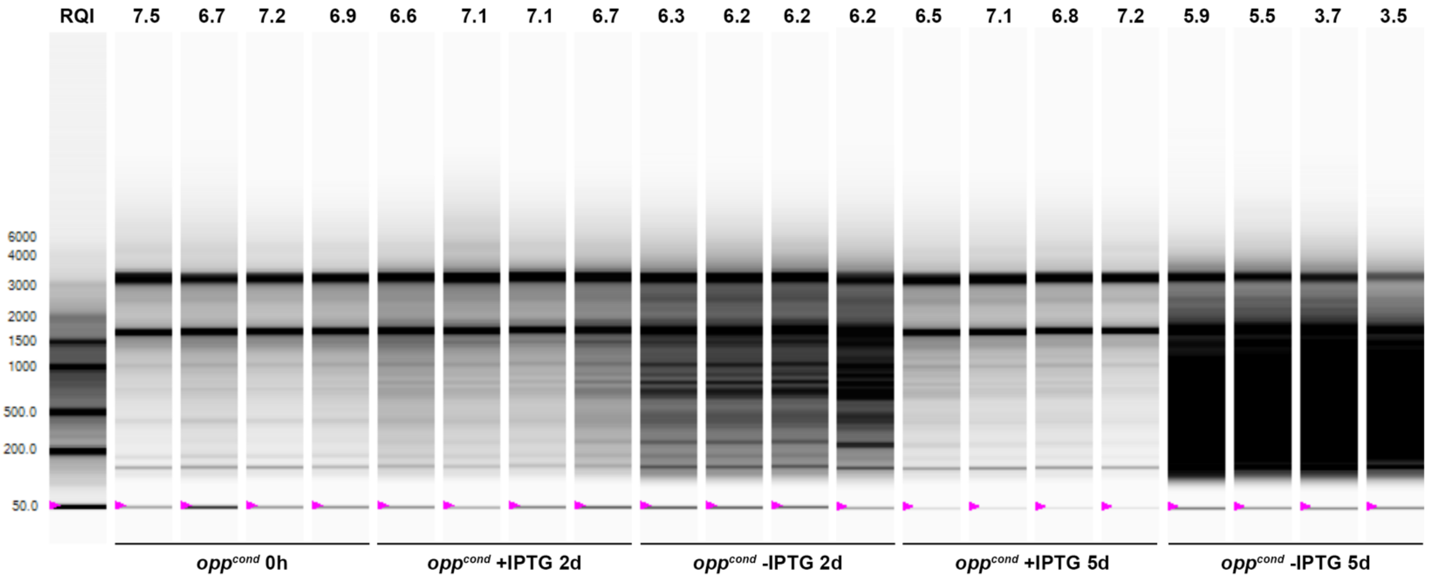


**Figure S6: Prolonged starvation of *opp*$^{cond}$ results in RNA degradation.** RNA samples collected from *opp*$^{cond}$ at timepoint 0 and without or with 1 mM IPTG at 2 d and 5 d post-incubation with their reported RNA quality indicator (RQI).

**Supplementary Table 1. *Borrelia burgdorferi* strains and plasmids used in this study**

| Strain/Plasmid | Description | Antibiotic Resistance | Reference |
|---|---|---|---|
| *B. burgdorferi* | | | |
| BbG100 | Wild-type strain B31 5A18 NP1 (*wt*) | Kan | [1] |
| BbG141A | B31 5A18 NP1 i*bb0334-35* Δ*bb0334-35* (*opp*$^{cond}$) | Kan/Strep/Erm | [2] |
| BbG141B | B31 5A18 NP1 i*bb0334-35* Δ*bb0334-35* (*opp*$^{cond+lp28\text{-}1}$) | Kan/Strep/Erm | This study |
| BbAG320 | B31 5A18 NP1 i*bb0401* (i*bb0401*) | Kan/Strep | This study |
| BbG119 | B31 5A18 NP1 i*bb0401* Δ*bb0401* (*bb0401*$^{cond}$) | Kan/Strep | This study |
| BbP1781 | B31 5A4 | N/A | [3] |
| BbAG367 | B31 5A4 i*gfp* (i*gfp*) | Strep | This study |
| *E. coli* | | | |
| Top10 | *F–mcrA Δ(mrr-hsdRMS-mcrBC) φ80lacZΔM15 ΔlacX74 recA1 araD139 Δ(ara-leu)7697 galU galK λ–rpsL(StrR) endA1 nupG* | N/A | Invitrogen |
| Stellar | *F–, endA1, supE44, thi-1, recA1, relA1, gyrA96, phoA, Φ80d lacZΔ M15, Δ (lacZYA - argF) U169, Δ (mrr - hsdRMS - mcrBC), ΔmcrA, λ–* | N/A | Clonetech |
| Plasmids | | | |
| pUC19 | Cloning vector | Amp | Invitrogen |
| pJSB275 | Shuttle vector with IPTG-inducible luciferase lacking NdeI in the resistance marker | Spec/Strep | [2] |
| pBRV2 | Gent marker | Gent | [4] |
| pEcAG286 | pi*bb0401* | Strep | This study |
| pEcAG259 | pΔ*bb0401* | Gent/Amp | This study |
| pCE320 | *gfp* | Zeo | [5] |
| EcAG304 | pi*gfp* | Strep | This study |

**Supplementary Table 2. Oligonucleotide primers used in this study**

| Designation | Sequence (5'-3') | Purpose | Reference |
|---|---|---|---|
| M13 F | CAGGAAACAGCTATGAC | Sequencing | Invitrogen |
| M13 R | GTAAAACGACGGCCAG | Sequencing | Invitrogen |
| pless Strep F | ATGAGGGAAGCGGTGATCGCCGA | Diagnostic PCR | [4] |
| pless Strep R | TTATTTGCCGACTACCTTGGTG | Diagnostic PCR | [4] |
| pless Gent F | ATGTTACGCAGCAGCAACGATG | Diagnostic PCR | [4] |
| pless Gent R | TTAGGTGGCGGTACTTGGGTCCA | Diagnostic PCR | [4] |
| 5' pJSB275 seq | GATTCAATTGTGAGCGGAATAACA | Sequencing | [6] |
| 3' pJSB275 seq | ATGCGCTTAACGGTAAAATCCAAGG | Sequencing | [6] |
| 5' bb0401 ind | **GGAGAAATTA*CATATG***AATATAAAAATCAATTTTTTTTTCACTTTG | i*bb0401*cloning | This study |
| 3' bb0401 ind | **CTCTATCTTC*AAGCTT***TTAATTAATTTTTTCTTGATCTTTTAATTCTTTG | i*bb0401*cloning | This study |
| 5' bb0401 null F1 | **CGACTCTAGA*GGATCC***GCCTCTTGGCCCTATC | Δ*bb0401* cloning | This study |
| 3' bb0401 null F1 | **TTGAAGCTCGGGTAG**ATGACTTCTCCTTTCAGAGATTTA | Δ*bb0401* cloning | This study |
| 5' bb0401 null gent | **GAAAGGAGAAGTCAT**CTACCCGAGCTTCAAGG | Δ*bb0401* cloning | This study |
| 3' bb0401 null gent | **TTAATTTGTTTAGCT**GGCGGTACTTGGGTC | Δ*bb0401* cloning | This study |
| 5' bb0401 null F2 | **GACCCAAGTACCGCC**AGCTAAACAAATTAATAGGATTTGGCA | Δ*bb0401* cloning | This study |
| 3' bb0401 null F2 | **CGGTACCCGG*GGATCC***GGCCTTTTTTGCGCAC | Δ*bb0401* cloning | This study |
| 5' iGFP | **AAGAGGAGAAATTA*CAT***_ATG_AGTAAAGGAGAAGAACTTTTC | i*gfp* cloning | This study |
| 3' iGFP | **CTCTATCTTC*AAGCTT***TTATTTGTATAGTTCATCCATGCC | i*gfp* cloning | This study |

Bold denotes overlap sequence for InFusion cloning. Italics denotes restriction sites.
